# Essential regulatory functions of CaMKII T286 phosphorylation in LTP and two distinct forms of LTD

**DOI:** 10.1101/2022.01.03.474759

**Authors:** Sarah G. Cook, Nicole L. Rumian, K. Ulrich Bayer

**Author notes:** Correspondence to: K.U.B.

## Abstract

The Ca^2+^/calmodulin-dependent protein kinase II (CaMKII) mediates both long-term potentiation and depression (LTP and LTD) of excitatory synapses, two opposing forms of synaptic plasticity induced by strong versus weak stimulation of NMDA-type glutamate receptors (NMDARs). NMDAR-dependent LTD is prevalent in juvenile hippocampus, but in mature hippocampus, LTD is still readily induced by stimulating metabotropic glutamate receptors (mGluRs). Here we show that mGluR-dependent LTD also requires CaMKII and its T286 autophosphorylation that induces Ca^2+^-independent autonomous kinase activity. This autophosphorylation (i) accelerated CaMKII movement to excitatory synapses after LTP stimuli and (ii) was required for the movement to inhibitory synapses after NMDAR-LTD stimuli. Similar to NMDAR-LTD, the mGluR-LTD stimuli did not induce any CaMKII movement to excitatory synapses. However, in contrast to NMDAR-LTD, the mGluR-LTD did not involve CaMKII movement to inhibitory synapses and did not require additional T305/306 autophosphorylation. Taken together, even though CaMKII T286 autophosphorylation has a longstanding prominent role in LTP, it is also required for both major forms of LTD in hippocampal neurons, albeit with differential requirements for the heterosynaptic communication of excitatory signals to inhibitory synapses.

## INTRODUCTION

CaMKII is a central mediator of NMDAR-dependent LTP and LTD (Bayer and Schulman, 2019), two opposing forms of synaptic plasticity thought to mediate higher brain functions such as learning, memory and cognition (Collingridge et al., 2010; Malenka and Bear, 2004; Martin et al., 2000). Both LTP and LTD require the CaMKII autophosphorylation at T286 that generates Ca^2+^-independent autonomous CaMKII activity (Coultrap et al., 2014; Giese et al., 1998). LTP additionally requires CaMKII binding to the NMDAR subunit GluN2B, which mediates the further accumulation of CaMKII at excitatory synapses (Barria and Malinow, 2005; Bayer et al., 2001; Halt et al., 2012; Incontro et al., 2018). By contrast, LTD instead requires additional inhibitory CaMKII autophosphorylation at T305/306 (Cook et al., 2021), which suppresses GluN2B binding and instead promotes CaMKII movement to inhibitory synapses, where it mediates inhibitory LTP (iLTP) (Barcomb et al., 2014; Cook et al., 2021). NMDAR-dependent LTD is still detectable in adult hippocampus but is much more prevalent at juvenile stages (Dudek and Bear, 1992, 1993). However, robust LTD can still be induced in mature hippocampus by stimulation of group 1 mGluRs, i.e. mGluR1 and 5 (Collingridge et al., 2010; Luscher and Huber, 2010; Reiner and Levitz, 2018). A role of CaMKII also in this mGluR-dependent LTD has been suggested by at least three independent pharmacological studies (Bernard et al., 2014; Mockett et al., 2011; Schnabel et al., 1999). However, these pharmacological studies differed in the direction of the reported effect. Thus, it remained unclear if CaMKII promotes or inhibits mGluR-dependent LTD.

Here, we tested CaMKII functions in mGluR-LTD using genetic approaches. Our results show that, like NMDAR-dependent LTP and LTD, the mGluR-LTD requires the CaMKIIα isoform and its autophosphorylation at T286. Thus, we additionally compared the role of T286 phosphorylation in the CaMKII targeting to excitatory versus inhibitory synapses in response to these three plasticity stimuli. In NMDAR-LTD, the function of T286 phosphorylation is to enable induction of additional T305/306 phosphorylation, which is known to be required for the CaMKII movement to inhibitory synapses and for normal NMDAR-LTD (Cook et al., 2021). By contrast, mGluR-LTD did not require this additional inhibitory autophosphorylation. Thus, even though CaMKII T286 autophosphorylation has a longstanding prominent role in LTP, it is also required for both major forms of LTD in hippocampal neurons, albeit with differential requirements for the heterosynaptic communication of excitatory signals to inhibitory synapses.

## RESULTS

### mGluR-LTD requires the CaMKIIα isoform

As pharmacological studies yielded conflicting results about the role of CaMKII in mGluR-LTD, we decided to test CaMKII functions in this form of plasticity by genetic means. Here, group I mGluRs (i.e. mGluR1 and mGluR5) were directly stimulated with DHPG (100 μM for 10 min). This treatment resulted in significant LTD at the CA3 to CA1 synapse in hippocampal slices from wild type mice (Figure 1A). Genetic knockout of the CaMKIIα isoform completely abolished this mGluR-LTD (Figure 1B). Thus, like LTP and NMDAR-LTD (Coultrap et al., 2014; Silva et al., 1992), normal mGluR-LTD specifically required the CaMKIIα isoform. Therefore, all additional CaMKII point mutations tested here were specifically in the α-isoform. Notably, the CaMKIIα isoform is exclusively expressed in neurons and is the major CaMKII isoform in mammalian brain, with ∼4fold higher hippocampal expression than the next most prevalent isoform, CaMKIIβ (Cook et al., 2018).

**Figure 1.**
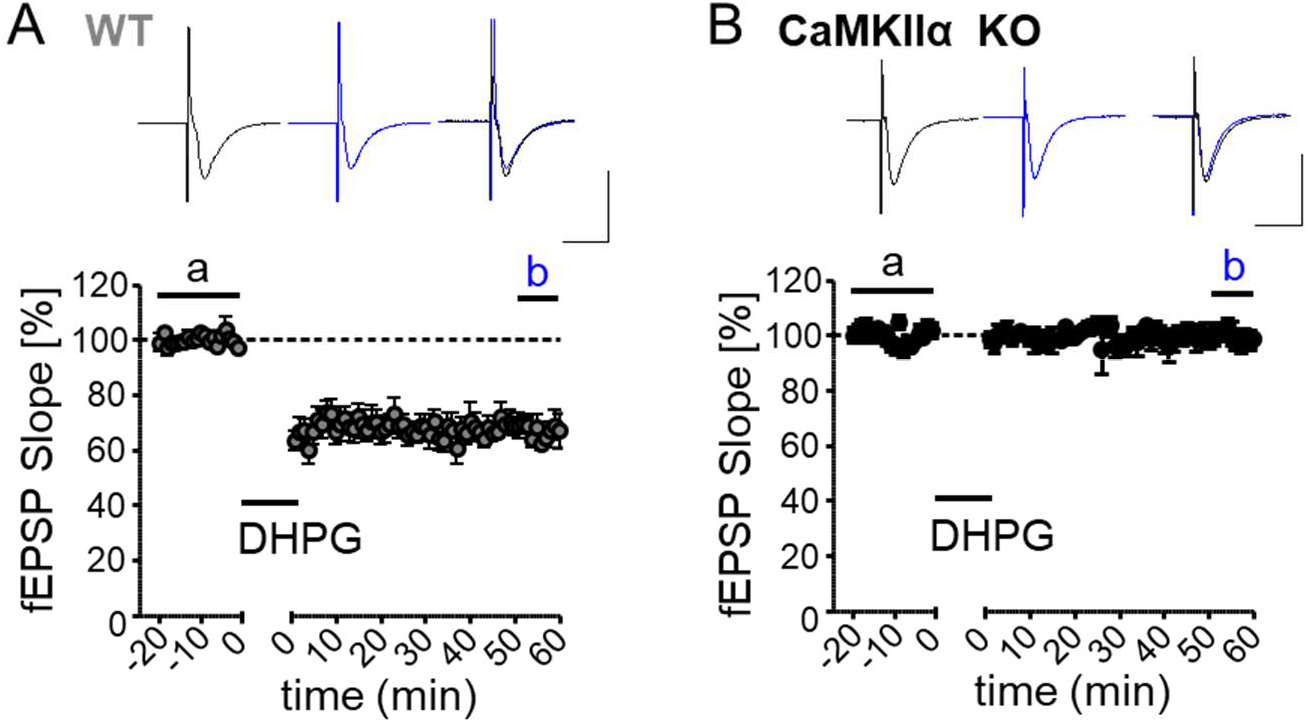
mGluR-LTD requires the CaMKIIα isoform. A Example traces and time course of synaptic response, measured by excitatory post-synaptic potential (EPSPs) slope of the CA3-CA1 Schaffer collateral pathway, in wild type hippocampal slices before and after chemical mGluR stimulation with DHPG (10 µM for 10 mins). B Example traces and time course of synaptic response in CaMKIIα KO slices before and after the same DHPG treatment as in panel A show abolished mGluR-LTD.

### mGluR-LTD requires CaMKII autophosphorylation at T286 but not T305/306

In order to determine the specific mechanisms of CaMKII regulation required for mGluR-LTD, we next tested the effect of several CaMKII point mutations. The T286A mutation prevents the T286 autophosphorylation that generates Ca^2+^-independent autonomous kinase activity (Colbran et al., 1989; Coultrap et al., 2010; Lou and Schulman, 1989; Miller and Kennedy, 1986). Like complete CaMKIIα knockout, this mutation completely abolished mGluR-LTD (Figure 2A). Thus, like LTP and NMDAR-LTD (Coultrap et al., 2014; Giese et al., 1998), mGluR-LTD specifically requires T286 autophosphorylation.

**Figure 2.**
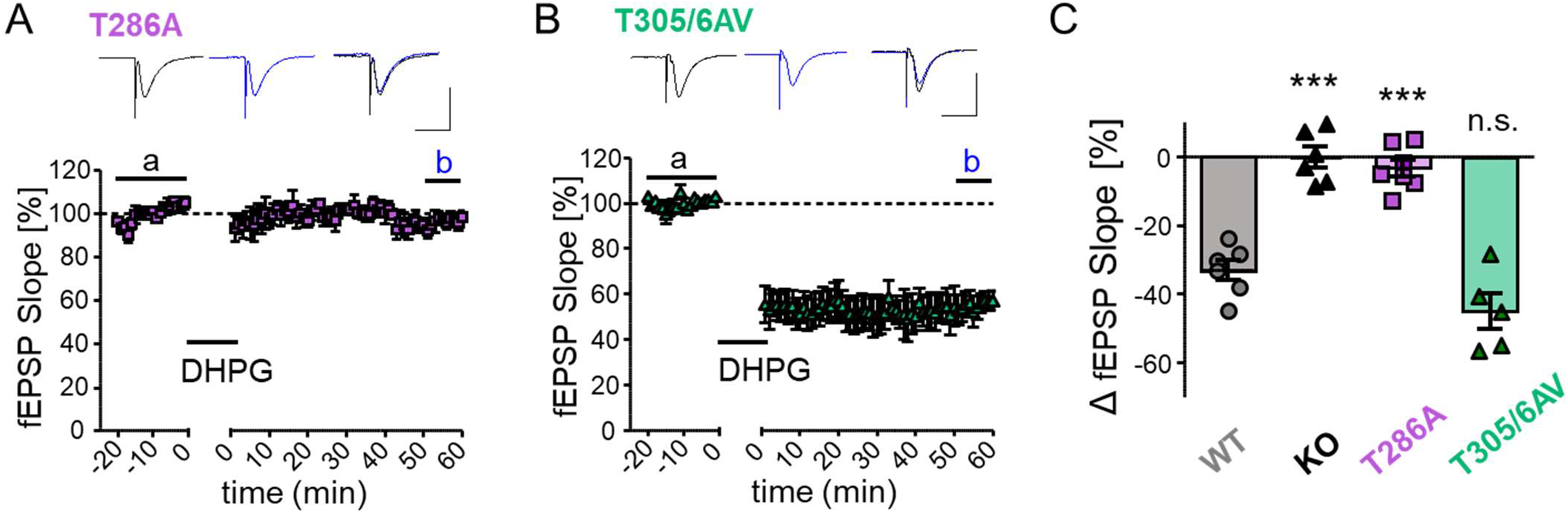
mGluR-LTD requires CaMKII auto-phosphorylation at T286 but not T305/306. A Example traces and time course of synaptic response in T286A slices before and after DHPG treatment. B Example traces and time course of synaptic response in T305/306AV slices before and after stimulation with DHPG (10 µM for 10 mins). C Quantification of the change in synaptic response (measured by EPSP slope) after DHPG stimulation in wild type, CaMKIIa KO, T286A, and T305/306AV slices. Both CaMKII KO and T286A animals demonstrated severe impairments in DHPG-induced mGluR LTD when compared to wild type animals while T305/306AV slices did not show any LTD deficit, indicating CaMKII and its T286 autophosphorylation are required for mGluR LTD (one-way ANOVA, Tukey’s post-hoc test vs. WT, ***p < 0.001, n=6, 6, 7, 5 slices).

An additional mutant tested was T305/306AV, which prevents an inhibitory autophosphorylation at these residues that blocks Ca^2+^/CaM-binding and also curbs autonomous kinase activity (Colbran and Soderling, 1990; Cook et al., 2021; Hanson and Schulman, 1992; Lou and Schulman, 1989). In contrast to the T286A mutation, the T305/306AV mutation did not reduce mGluR-LTD at all (Figure 2B). If any, the mGluR-LTD in the T305/306AV mice appeared to be slightly enhanced compared to wild type, however, this apparent enhancement was not statistically significant (Figure 2C). Thus, in contrast to NMDAR-LTD (Cook et al., 2021) but like LTP (Elgersma et al., 2002), mGluR-LTD does not require the inhibitory autophosphorylation at T305/306.

Figure 2C summarizes the effects on mGluR-LTD of all CaMKII mutant mice tested here. Overall, CaMKIIα and its T286 autophosphorylation is required for normal LTP, for NMDAR-LTD, and for mGluR-LTD. By contrast, additional T305/306 autophosphorylation is required only for normal NMDAR-LTD, but not for LTP or mGluR-LTD.

### T286 autophosphorylation accelerates movement of endogenous CaMKII to excitatory synapses after LTP stimuli

As T286 autophosphorylation is required for three distinct forms of long-term synaptic plasticity, we decided to determine how it affects CaMKII movement in hippocampal neurons in response to the different plasticity stimuli. It is known that T286 autophosphorylation is not strictly required for CaMKII movement to excitatory synapses in response to chemical LTP (cLTP) stimuli (Shen and Meyer, 1999) or for Ca^2+^/CaM-induced binding to GluN2B *in vitro* (Bayer et al., 2001). Indeed, the CaMKII movement induced by cLTP stimuli (1 min 100 μM glutamate in the presence of 10 µM glycine) was indistinguishable between overexpressed GFP-CaMKII wild type and its T286A mutant (Figure 3). However, when we instead monitored the movement of endogenous CaMKII with our intrabody method (Cook et al., 2021; Cook et al., 2019), CaMKII movement was significantly faster in wild type neurons compared to neurons from T286A mutant mice (Figure 4). Thus, even though T286 autophosphorylation is not essential for the CaMKII movement to excitatory synapses that is required for normal LTP (Barria and Malinow, 2005; Halt et al., 2012), it significantly accelerates the process.

**Figure 3.**
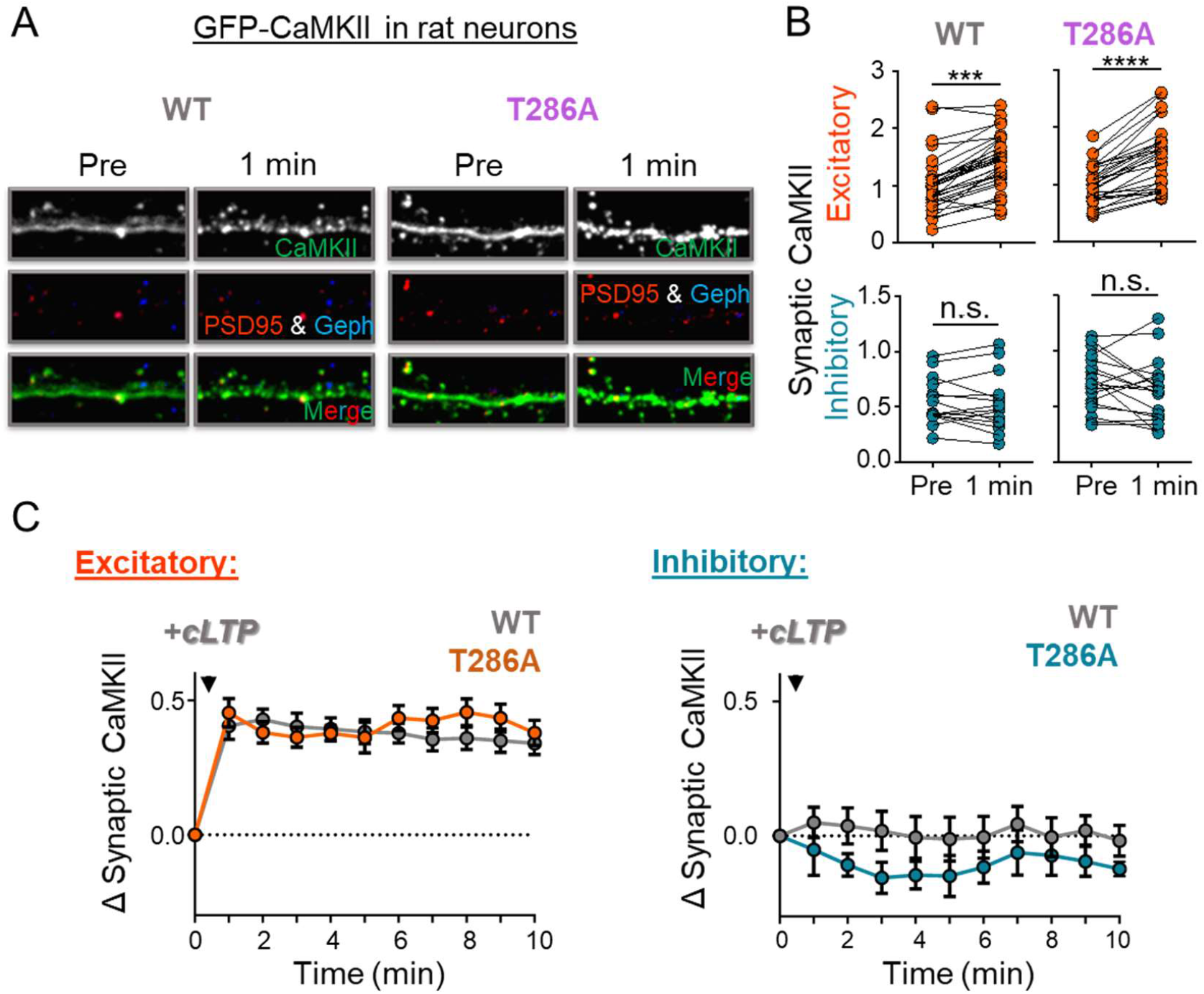
T286A mutation does not affect the LTP-induced movement of overexpressed GFP-CaMKII. A Representative images of rat hippocampal neurons (DIV 14-17) expressing intrabodies for detection of PSD-95 (red) and gephyrin (blue) to label excitatory and inhibitory synapses, respectively, and overexpressing either CaMKII wild type or T286A (green) before and 1 min following chemical NMDAR-LTP stimulation (cLTP; 100 µM glutamate/10 µM glycine, 1 min). B Quantification of wild type or T286A CaMKII at excitatory (red) and inhibitory (blue) synapses before and 1 min post cLTP stimulation. Both overexpressed wild type and T286A CaMKII translocated to excitatory, but not inhibitory, synapses following cLTP treatment (paired t-test, wild type, left panels: ****p < 0.0001, T286A, right panels: ***p = 0.0003, n = 21, 12 neurons). C Full time course of wild type (grey) and T286A CaMKII (orange) movement to excitatory and inhibitory synapses following cLTP stimulation indicating similar translocation dynamics for both constructs.

**Figure 4.**
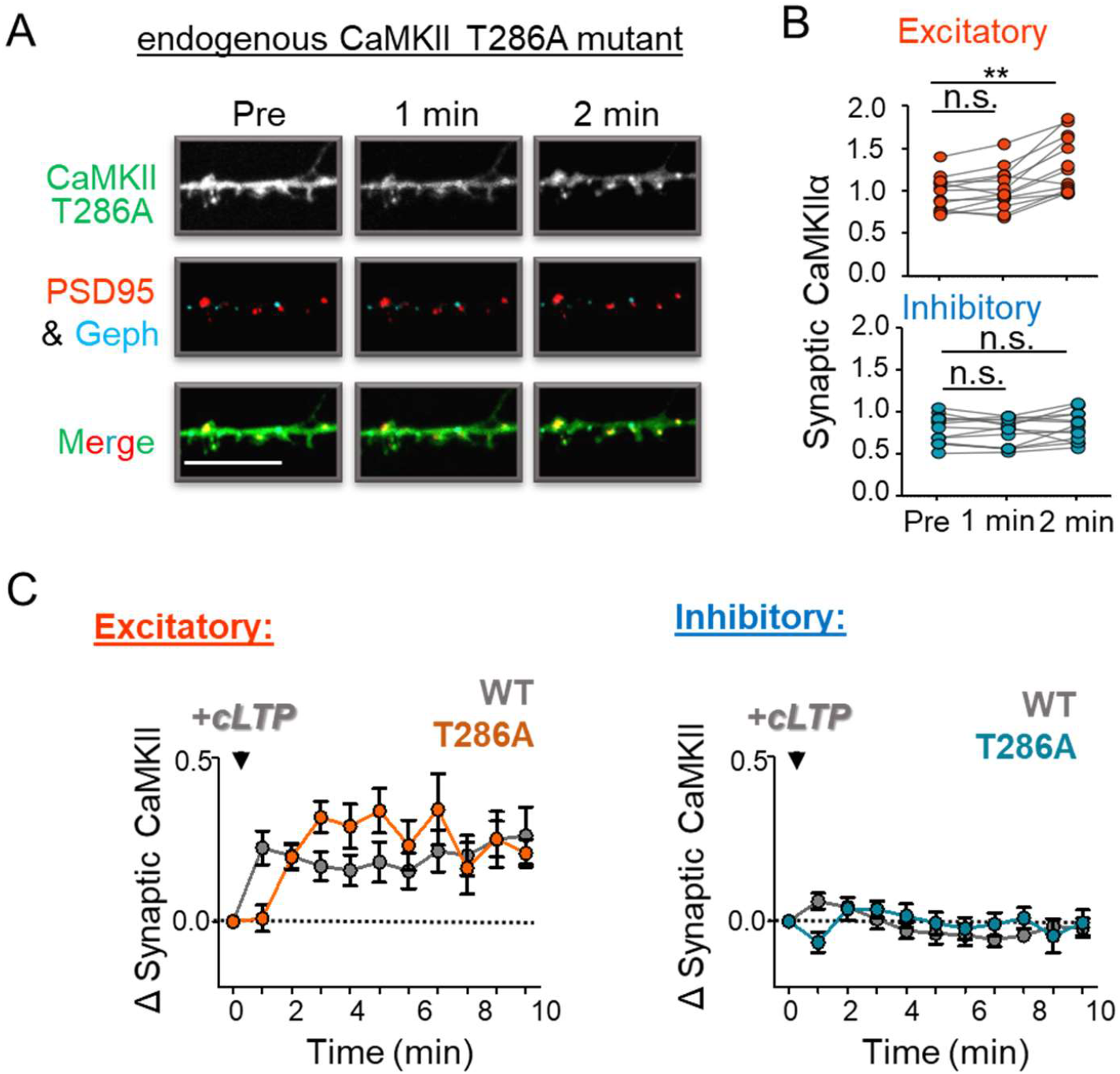
LTP-induced movement of endogenous CaMKII to excitatory synapses is accelerated by T286 autophosphorylation. A Representative images of hippocampal neurons from CaMKII T286A mutant mice (DIV 14-17) expressing intrabodies against endogenous PSD-95 (red) and gephyrin (blue), to label excitatory and inhibitory synapses, respectively, and CaMKII (green) before, 1 min, and 2 min after cLTP treatment. Scale bar in the example images indicates 10 µM. B Quantification of CaMKII T286A at excitatory (red) and inhibitory (blue) synapses pre, 1 min, and 2 min post cLTP stimulation. Similar as described in WT neurons, endogenous CaMKII in hippocampal neurons from T286A mutant mice moved to excitatory but not inhibitory synapses in response to cLTP stimulation. However, the endogenous T286A mutant moved more slowly to excitatory synapses, as significant synaptic enrichment was seen only at 2 min but not at 1 min after cLTP stimuli. These results indicate T286 autophosphorylation accelerates LTP-induced CaMKII synaptic targeting (one-way ANOVA, Tukey’s post-hoc test vs. pre, **p = 0.0054, n = 13 neurons). C Full time course of T286A CaMKII movement to excitatory and inhibitory synapses following cLTP stimulation. For comparison, the movement of wild type CaMKII is illustrated in grey; this movement was measured in parallel experiment but was published previously (Cook et al., 2021).

CaMKII movement to inhibitory synapses was not detected after cLTP stimuli, neither for overexpressed nor for endogenous CaMKII (see Figures 3 and 4), as expected based on our previous results (Cook et al., 2021; Cook et al., 2019).

### T286 autophosphorylation is required for CaMKII movement to inhibitory synapses after NMDAR-LTD stimuli

In contrast to cLTP stimuli, NMDAR-dependent chemical LTD (cLTD) stimuli (30 μM NMDA, 10 μM glycine, and 10 μM CNQX for 1 min) cause CaMKII movement to inhibitory synapses, and this movement strictly requires T305/306 autophosphorylation (Cook et al., 2021). An additional co-requirement of T286 phosphorylation should be expected, because it is required for efficient T305/306 phosphorylation (Cook et al., 2021). Indeed, such requirement for T286 phosphorylation has been suggested by experiments with overexpressed GFP-CaMKII (Marsden et al., 2010). Nonetheless, due to the discrepancies between movement of overexpressed and endogenous CaMKII in response to cLTP stimuli (see Figures 3 and 4), we decided to examine the effect of T286A mutation also on endogenous CaMKII. As predicted, the T286A mutation completely abolished CaMKII movement to inhibitory synapses in response to NMDAR-dependent cLTD stimuli (Figure 5A-C). Thus, whereas T286 autophosphorylation affects CaMKII movement to excitatory synapses in its temporal aspects, it is absolutely required for any CaMKII movement to inhibitory synapses.

**Figure 5.**
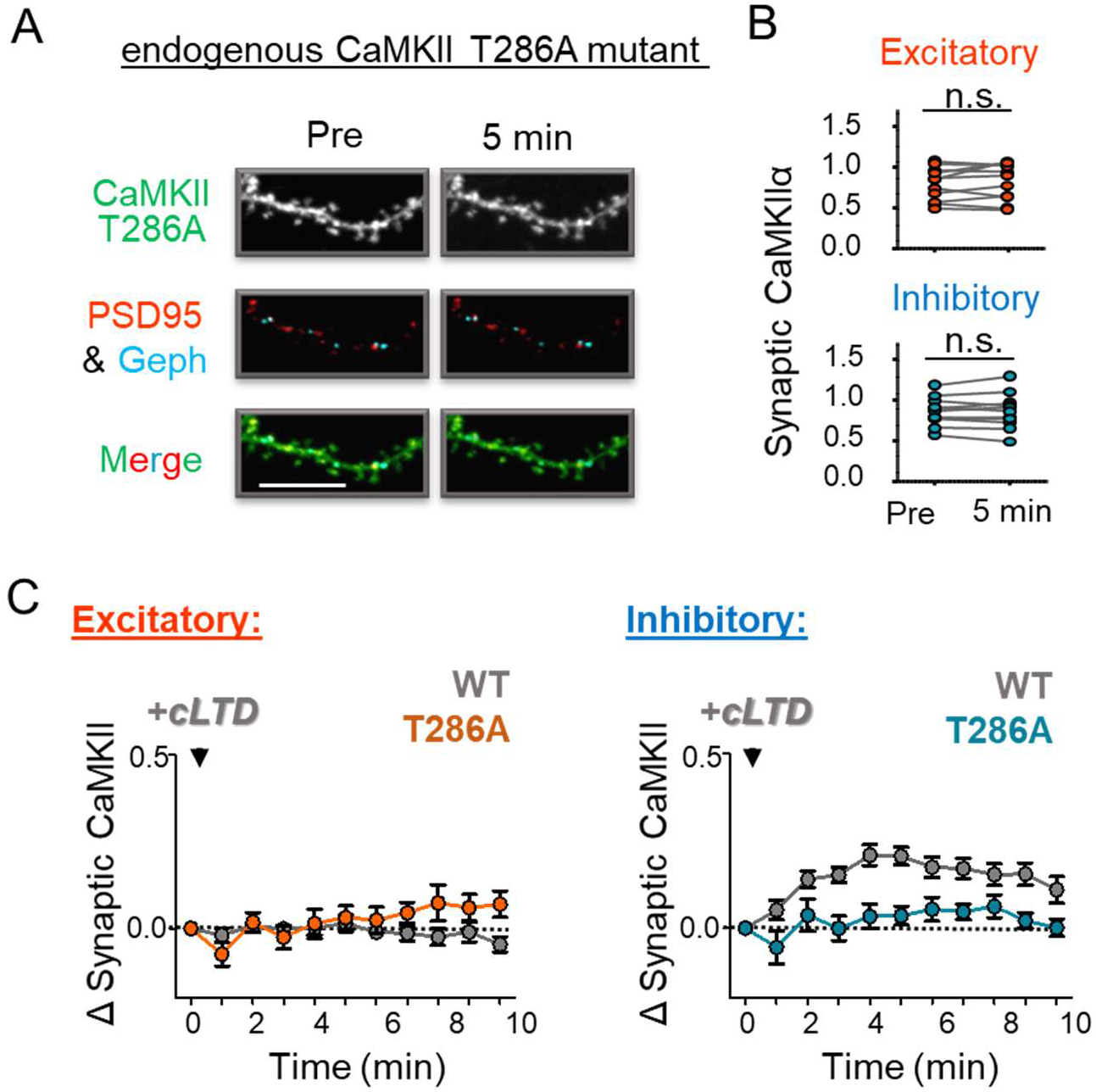
CaMKII movement to inhibitory synapses in response to NMDAR-LTD stimuli requires T286 autophosphorylation. A Representative images of hippocampal neurons from CaMKII T286A mutant mice (DIV 14-17) expressing intrabodies against endogenous PSD-95 (red) and gephyrin (blue), to label excitatory and inhibitory synapses, respectively, and CaMKII (green) before and 5 min after chemical NMDAR-LTD stimuli (cLTD; 30 µM NMDA/10 µM CNQX/10 µM glycine, 1 min). Scale bar in the example images indicates 10 µM. B Quantification of T286A CaMKII at excitatory (red) and inhibitory (blue) synapses prior to and 5 min post cLTD stimulation. In response to cLTD, endogenous T286A mutant CaMKII did not move to either excitatory or inhibitory synapses (in contrast to CaMKII wild type, which has been described to move to inhibitory but not excitatory synapses in response to LTD). C Lack of movement of the T286A mutant to either excitatory (red) or inhibitory (blue) synapses following cLTD is further illustrated in a full time course. For comparison, the movement of wild type CaMKII is illustrated in grey; this movement was measured in parallel experiment but was published previously (Cook et al., 2021).

### mGluR-LTD stimuli do not induce synaptic CaMKII movement

As NMDAR-dependent cLTP or cLTD stimuli induce CaMKII movement to either excitatory or inhibitory synapses, respectively, we decided to examine if mGluR-LTD stimuli with 100 μM DHPG for 10 min also cause CaMKII movement. Endogenous CaMKII was monitored for 20 min after the DHPG stimulus in cultured hippocampal neurons, but no movement to either excitatory or inhibitory synapses was detected (Figure 6A). After NMDAR-dependent cLTP stimuli, lack of CaMKII movement to excitatory synapses is mediated by active suppression mechanisms (Cook et al., 2021; Goodell et al., 2017), and the CaMKII T305/306AV mutation is sufficient to restore movement to excitatory synapses even after such cLTD stimuli (Cook et al., 2021). Thus, we also tested neurons from T305/306AV mutant mice for DHPG-induced CaMKII movement, but again, no movement was detected (Figure 6B). This indicates either that the mechanisms for suppression of CaMKII movement differ in NMDAR-versus mGluR-LTD or that mGluR-LTD may not require such a suppression mechanism at all. Either scenario is consistent with the normal mGluR-LTD observed in hippocampal slices from the CaMKII T305/T306AV mutant mice (see Figure 2B,C).

**Figure 6.**
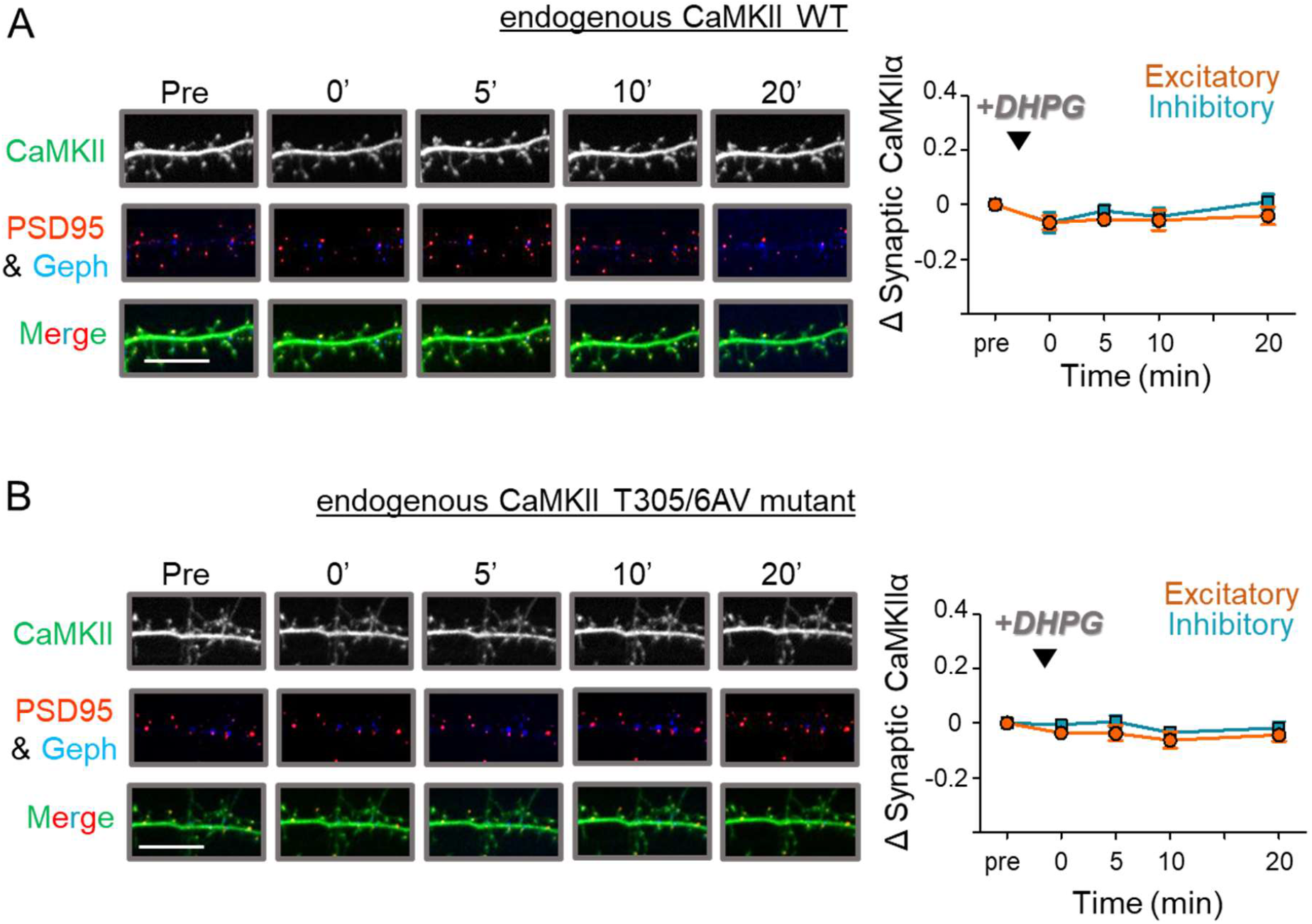
mGluR stimulation does not promote CaMKII movement to synapses. A Representative images of rat hippocampal neurons (DIV 14-17) expressing intrabodies targeting endogenous CaMKII (green), as well as PSD-95 (red) and gephyrin (blue) to label excitatory and inhibitory synapses respectively, before and 0, 5, 10, and 20 min after stimulation with DHPG (100 μM for 10 min). Scale bar in the example images indicates 10 µM. The left panel shows a full time course quantification of endogenous wild type CaMKII at excitatory (red) and inhibitory (blue) synapses prior to and 0, 5, 10, and 20 min post DHPG stimulation. There was no CaMKII movement to either excitatory or inhibitory synapses was found in response to DHPG stimulation (in contrast to NMDAR-LTD stimuli, where wild type CaMKII has been described to move to inhibitory but not excitatory synapses). B Representative images of hippocampal neurons from CaMKII T305/306AV mutant mice (DIV 14-17) expressing intrabodies targeting endogenous CaMKII (green), as well as PSD-95 (red) and gephyrin (blue) to label excitatory and inhibitory synapses respectively, before and 0, 5, 10, and 20 min after stimulation with DHPG (100 μM for 10 min). Scale bar in the example images indicates 10 µM. The left panel shows a full time course quantification of endogenous T305/6AV CaMKII at excitatory (red) and inhibitory (blue) synapses prior to and 0, 5, 10, and 20 min post DHPG stimulation. Again, no movement of CaMKII to either excitatory or inhibitory synapses was found in response to DHPG stimulation.

Thus, overall, even though DHPG-induced mGluR-LTD does not induce any synaptic CaMKII movement, it does require the CaMKIIα isoform and its T286 phosphorylation (which accelerates the CaMKII movement to excitatory synapses after LTP and is absolutely required for the CaMKII movement to inhibitory synapses after NMDAR-LTD), but not its T305/306 phosphorylation (which is required for the specific suppression of CaMKII movement to excitatory synapses during NMDAR-LTD).

## DISCUSSION

CaMKII is a central regulator of synaptic plasticity and well established to mediate both NMDAR-dependent LTP and LTD (for review see (Bayer and Schulman, 2019). Here, we demonstrate an additional requirement for CaMKII also in mGluR-LTD. Like NMDAR-LTP, but in contrast to NMDAR-LTD, this mGluR-LTD is robustly induced not only in young but also in mature animals. Like both forms of NMDAR-dependent plasticity (Coultrap et al., 2014; Giese et al., 1998; Silva et al., 1992), the mGluR-LTD required the CaMKIIα isoform and its autophosphorylation at T286. However, in contrast to NMDAR-LTD (Cook et al., 2021), the mGluR-LTD did not require additional autophosphorylation at T305/306. Consistent with this electrophysiological observation in hippocampal slices, imaging in hippocampal neurons showed that mGluR-LTD did not require T305/306 phosphorylation for suppression of CaMKII movement to excitatory synapses. This CaMKII movement is required for normal NMDAR-LTP (Barria and Malinow, 2005; Halt et al., 2012) and has to be actively suppressed during NMDAR-LTD (Cook et al., 2021; Goodell et al., 2017). If any active suppression of CaMKII movement is also required for mGluR-LTD, the mechanism must differ from NMDAR-LTD, as the latter requires T305/306 phosphorylation (Cook et al., 2021) whereas our results show that the former does not.

T286 phosphorylation is required for all three forms of long-term synaptic plasticity, but the time course of phosphorylation appears to differ. Phosphorylation at T286 generates Ca^2+^-independent “autonomous” activity and LTP stimuli had been proposed to cause a self-perpetuated increase in T286 phosphorylation. However, T286 autophosphorylation occurs between two different subunits of the 12meric CaMKII holoenzyme and requires Ca^2+^/CaM binding to both subunits (Hanson et al., 1994; Rich and Schulman, 1998), and this mechanism does not support the originally proposed perpetuation mechanism. Additionally, imaging experiments indicated that while T286 autophosphorylation indeed extends the CaMKII activation state after LTP-stimuli, this was only on a time scale of less than 2 min (Lee et al., 2009). Nonetheless, the fast reversal of T286 phosphorylation after LTP remained somewhat controversial, in part because the imaging experiments did not directly assess T286 phosphorylation (for review see (Lisman et al., 2012). However, one of our recent studies directly compared T286 phosphorylation after NMDAR-dependent LTP versus LTD, and indicated that LTP stimuli caused a larger increase in T286 phosphorylation, but that this increase was fully reversed within 5 min (Cook et al., 2021). After NMDAR-LTD, the increase in T286 phosphorylation also appeared to be immediate but was then maintained longer (Cook et al., 2021). By contrast, the mGluR-LTD stimulus caused a delayed increase in T286 phosphorylation that was not apparent immediately but was significant after 5 min (Mockett et al., 2011).

The different temporal pattern of T286 phosphorylation may help to enable the opposing downstream effects of CaMKII in the different forms of long-term plasticity. LTP additionally requires CaMKII binding to GluN2B (Barria and Malinow, 2005; Halt et al., 2012), whereas NMDAR-LTD instead additionally requires T305/306 autophosphorylation (Cook et al., 2021). Notably, GluN2B binding and T305/306 phosphorylation are both promoted by T286 phosphorylation (Bayer et al., 2001; Strack et al., 2000) but then mutually inhibit each other (Barcomb et al., 2014; Bayer et al., 2001). Thus, the different extent and time course of T286 phosphorylation may decide which one of the two downstream events after T286 phosphorylation prevails, i.e. the GluN2B binding that is required for normal LTP or the T305/306 phosphorylation that is required for normal NMDAR-LTD. The mGluR-LTD appears to require neither of these two mechanisms: mGluR-LTD did not cause a GluN2B-mediated CaMKII accumulation in hippocampal neuron, and CaMKII phosphorylation at T305/306 was also not required, neither for the suppression of movement nor for the mGluR-LTD detected in hippocampal slices. While mGluR-LTD was expected to not require the GluN2B binding, lack of requirement of pT305/306 was somewhat more surprising. In NMDAR-LTD, such additional T305/306 phosphorylation is required to suppress CaMKII movement to excitatory synapses and instead direct CaMKII movement to inhibitory synapses, where it then induces inhibitory LTP (Cook et al., 2021). Consistent with a lack of T305/306 phosphorylation after mGluR-LTD, no CaMKII movement to inhibitory synapses was observed during this form of plasticity. Thus, whereas mGluR- and NMDAR-LTD both decrease the strength of excitatory synapses, only NMDAR-LTD stimuli appear to elicit additional heterosynaptic communication to inhibitory synapses.

The Ca^2+^/CaM-induced CaMKII binding to GluN2B appears to be both necessary and sufficient for the further accumulation of CaMKII at excitatory synapses during LTP (Bayer et al., 2001; Halt et al., 2012). However, CaMKII can bind to various other synaptic proteins (Bayer and Schulman, 2001; Colbran, 2004; Hell, 2014; Kim et al., 2016), including mGluR1 and mGluR5. Interestingly, Ca^2+^/CaM stimulates CaMKII binding to mGluR1 but disrupts binding to mGluR5, whereas CaMKII T286 autophosphorylation may increase binding to both (Jin et al., 2013a; Jin et al., 2013b; Marks et al., 2018). CaMKII binding and phosphorylation can regulate these mGluRs, but any potential contribution to the mGluR-LTD studied here is currently unknown. Notably, even though normal NMDAR-LTP requires CaMKII binding to GluN2B, the opposing NMDAR-LTD does not (Goodell et al., 2017; Halt et al., 2012). Furthermore, mGluR-LTD stimuli did not induce any detectable synaptic CaMKII movement. These parallels and observation do not rule out a function of CaMKII binding to mGluRs in mGluR-LTD. However, a direct function of CaMKII/mGluR binding in mGluR-LTD seems less conceivable than a more indirect function in mGluR metaplasticity, i.e. in mediating signaling that modulates induction of subsequent mGluR-LTD. It will be interesting to elucidate the possible involvement of CaMKII protein-protein interactions in regulating different functions of mGluR-mediated plasticity, based on the requirement of CaMKII and its autophosphorylation at T286 but not T305/306 that was demonstrated here.

## METHODS

### Animals

All animal procedures were approved by the University of Colorado Institutional Animal Care and Use Committee (IACUC) and carried out in accordance with NIH best practices for animal use. The University of Colorado Anschutz Medical Campus is accredited by the Association for Assessment and Accreditation of Laboratory Animal Care, International (AAALAC). All animals were housed in ventilated cages on a 12 h light/ 12 h dark cycle and were provided ad libitum access to food and water. Mixed sex WT or mutant mouse littermates (on a C57BL/6 background) from heterozygous breeder pairs (8-12 weeks old) were used for slice electrophysiology and biochemistry. Mixed sex pups from homozygous mice (P1-2) or Sprague-Dawley rats (P0, Charles River) were used to prepare dissociated hippocampal cultures for imaging. CaMKIIα knock-out, T286A, and T305/306AV mice were described previously (Coultrap et al., 2014; Elgersma et al., 2002; Giese et al., 1998).

### Mouse hippocampal slice preparation

Isoflurane anesthetized mice were rapidly decapitated, and the brain was dissected in ice-cold high sucrose solution containing (in mM): 220 sucrose, 12 MgSO4, 10 glucose, 0.2 CaCl2, 0.5 KCl, 0.65 NaH2PO4, 13 NaHCO3, and 1.8 ascorbate. Whole and CA1 mini hippocampal slices (400 µm) were made using a tissue chopper (McIlwain) with CA1 mini-slice preparation requiring additional cuts as described previously (Cook et al., 2021). Slices were transferred into 32°C artificial cerebral spinal fluid (ACSF) containing (in mM): 124 NaCl, 2 KCl, 1.3 NaH2 PO4, 26 NaHCO3, 10 glucose, 2 CaCl2, 1 MgSO4, and 1.8 ascorbate and recovered in 95% O2/5% CO2 for at least 1.5 h before experimentation.

### Primary rat and mouse hippocampal culture

Primary hippocampal neurons were cultured as described previously (Vest et al., 2010) and imaged after 14-17 days in vitro (DIV14-17). Pups were decapitated and hippocampi were dissected and incubated in dissociation solution (7 mL HBSS buffered saline, 150 μL 100mM CaCl2, 10 μL 1M NaOH, 10 μL 500mM EDTA, 200 units Papain [Worthington]) at 25°C for 1 h (rat) or 30 mins (mice). Hippocampi were then washed 5x with plating media (DMEM, FBS, 50 units/ml Penn/strep, 2 mM L-glutamine, filter sterilized) and manually dissociated and counted using a hemocytometer. Dissociated neurons were plated on poly-D-lysine (0.1mg/mL in 1M Borate Buffer: 3.1g Boric Acid, 4.75 g borax, in 1 L deionized H20, filter sterilized) and laminin (0.01mg/mL in PBS)-coated 18 mm glass coverslips in 12 well plates at a density of 75,000-100,000 (rat) or 150,000-200,000 (mice) neurons per well in plating media and maintained at 37°C with 5% CO2. After 1 day in vitro (DIV 1), media was switched to 100% feeding media (Neurobasal-A, B27 supplements, and 2 mM L-glutamine, filter sterilized). At DIV At DIV 5, 50% of media was replaced with fresh neuron feeding media and treated with FDU (70 μM 5-fluoro-2′-deoxyuridine/140 μM uridine) to suppress glial growth by halting mitosis. At 12-14 DIV, neurons were transfected with the intrabodies using a 1:1:1 ratio, at a concentration of 1 μg total DNA/well, using Lipofectamine 2000 (2.5 μL/well, Invitrogen) according to the manufacturer’s recommendations.

### Extracellular field recordings

All recordings and analysis were performed blind to genotype. For electrical slice recording experiments, a glass micro-pipette (typical resistance 0.4 to 0.8 MΩ when filled with ACSF) was used to record field excitatory post-synaptic potentials (fEPSPs) from the CA1 dendritic layer in response to stimulation in the Schaffer collaterals at the CA2 to CA1 interface using a tungsten bi-polar electrode. Slices were continually perfused with 30.5 ± 0.5°C ACSF at a rate of 3.5 ± 0.5 mL/min during recordings. Stimuli were delivered every 20 sec and 3 responses (1 min) were averaged for analysis. Data were analyzed using WIN LTP (Anderson and Collingridge, 2001) with slope calculated as the initial rise from 10 to 60% of response peak. Input/output (I/O) curves were generated by increasing the stimulus intensity at a constant interval until a maximum response or population spike was noted to determine stimulation that elicits 40-70% of maximum slope. Slope of I/O curve was calculated by dividing the slope of response (mV/ms) by the fiber volley amplitude (mV) for the initial linear increase. Paired-pulse recordings (50 ms inter-pulse interval) were acquired from 40% max slope and no differences in presynaptic facilitation were seen in mutant slices. A stable baseline was acquired for a minimum of 20 min at 70% maximum slope prior to mGluR-LTD induction using 100 μM DHPG ((S)-3,5-Dihydroxyphenylglycine) for 10 mins. Slices were perfused with for the remainder of the recording in ACSF containing an NMDAR-antagonist (50 µM APV). Change in slope was calculated as a ratio of the average slope of the 20 min baseline (prior to stimulation).

### Chemical LTP and LTD stimulation

NMDAR-dependent LTP (NMDAR-LTP) was chemically induced using 100 µM glutamate and 10 μM glycine for 1 min. NMDAR-dependent LTD (NMDAR-LTD) was chemically induced with 30 μM NMDA, 10 μM glycine, and 10 μM CNQX for 1 min. mGluR-dependent LTD (mGluR-LTD) was induced with 100 μM DHPG for 10 min. All treatments were followed by 5X washout in fresh ACSF. For imaging and biochemical experiments, quantifications demonstrate the change from 1 min pre stimulation and 1 min (cLTP) or 5 mins (cLTD) post-wash out unless otherwise specified.

### Imaging Acquisition and Analysis

All microscopic imaging was performed using a 100 × 1.4NA objective on a Zeiss Axiovert 200 M (Carl Zeiss, Thornwood, NY) controlled by SlideBook software (Intelligent Imaging Innovations, Denver, CO). All imaging analysis was completed using SlideBook software. All representative images were prepared using Fiji software (Imagej, NIH). For all imaging experiments, focal plane z stacks (0.3-µm steps; over 1.8–2.4 µm) were acquired and deconvolved to reduce out-of-focus light. 2D maximum intensity projection images were then generated and analyzed by an experimenter blinded to experimental conditions. During image acquisition, neurons were maintained at 34°C in ACSF solution containing (in mM): 130 NaCl, 5 KCl, 10 HEPES pH 7.4, 20 glucose, 2 CaCl2, and 1 MgCl2, adjusted to proper osmolarity with sucrose. After baseline imaging and cLTP or cLTD treatment, neurons were imaged once per minute for 10 mins to limit the effects of photobleaching.

Hippocampal neurons were selected based on pyramidal shaped soma and presence of spiny apical dendrites, and tertiary dendritic branches were selected for analysis to maintain consistency. Images were analyzed at 1 min before stimulation and 1 min (after cLTP), 5 mins (after cLTD), and 10 mins (after DHPG) after wash out. 2D maximum intensity projection images were then generated and analyzed by an experimenter blinded to condition using Slidebook 6.0 software. To analyze endogenous YFP-intrabody-labelled CaMKII and overexpressed GFP-labelled CaMKII, the mean YFP or GFP intensity (CaMKII) at excitatory (PSD-95) and inhibitory (gephyrin) synapses was quantified. PSD-95 and gephyrin threshold masks were defined using the mean intensity of mCh or mTurquois plus two standard deviations. Synaptic CaMKII was then calculated using the mean YFP or GFP intensity at PSD-95 or gephyrin puncta masks divided by the mean intensity of a line drawn in the dendritic shaft. Changes in CaMKII synaptic accumulation were determined by dividing the net change in CaMKII at PSD-95 or gephyrin puncta-to shaft ratio by the pre-stimulation puncta-to-shaft ratio.

### Statistical analysis

Data are shown as mean ± SEM and all imaging and western blot quantification is normalized to control average set to 1. Statistical significance and sample size (n) are indicated in the Figure legends. Data obtained from imaging experiments were obtained using SlideBook 6.0 software (3i) and analyzed using Prism (GraphPad) software. All data met parametric conditions, as evaluated by a Shapiro-Wilk test for normal distribution and a Brown-Forsythe test (3 or more groups) or an F-test (2 groups) to determine equal variance. Comparisons between two groups were analyzed using unpaired, two-way Student’s t tests. Comparisons between pre- and post-treatment images from the same cells were analyzed using paired, two-way Student’s t tests. Comparisons between three or more groups were done by one-way ANOVA with Tukey’s post hoc analysis. Comparisons between three or more groups with two independent variables were accessed using a two-way ANOVA to determine whether there is an interaction and/or main effect between the variables. Statistical significance is indicated, including by *p < 0.05; **p < 0.01; ***p < 0.001, **** p<0.0001.

## ACKNOWLEDGMENTS

This research was supported by National Institutes of Health grants F31AG066536 (to S.G.C.), F31AG069458 (to N.L.R.), R01NS081248 and R01AG067713 (to K.U.B.).

## AUTHOR CONTRIBUTIONS

S.G.C. and N.L.R. performed experiments and analysis. S.B.C and K.U.B. conceived the study. K.U.B. wrote the first draft with input from all authors.

## CONFLICT OF INTEREST STATEMENT

K.U.B. is co-founder and board member of Neurexis Therapeutics.

